# Systems-level physiology of the human red blood cell is computed from metabolic and macromolecular mechanisms

**DOI:** 10.1101/797258

**Authors:** James T. Yurkovich, Laurence Yang, Bernhard O. Palsson

## Abstract

The human red blood cell has served as a starting point for the application and development of systems biology approaches due to its simplicity, intrinsic experimental accessibility, and importance in human health applications. Here, we present a multi-scale computational model of the human red blood cell that accounts for the full metabolic network, key proteins (>95% of proteome mass fraction), and several macromolecular mechanisms. Proteomics data are used to place quantitative constraints on individual protein complexes that catalyze metabolic reactions, as well as a total proteome capacity constraint. We explicitly describe molecular mechanisms—such as hemoglobin binding and the formation and detoxification of reactive oxygen species—and takes standard hematological variables (e.g., hematocrit, hemoglobin concentration) as input, allowing for personalized physiological predictions. This model is built from first principles and allows for direct computation of physiologically meaningful quantities such as the oxygen dissociation curve and an accurate computation of the flux state of the metabolic network. More broadly, this work represents an important step toward including the proteome and its function in whole-cell models of human cells.

## INTRODUCTION

The blood is a window through which we can explore human health and disease [50]. The human red blood cell (RBC)—the most abundant human cell [46]—has historically been used as the starting point for the application and development of systems biology models due to its relative simplicity, intrinsic accessibility, and the vast amounts of data and information available on its biochemistry and physiology [52]. While the RBC lacks cellular compartments and the ability to produce energy using oxidative phosphorylation, it is heavily involved in the transport and exchange of gases throughout the body, including O_2_, CO_2_ [17], and NO [15]. The ability to model and compute physiological states of the RBC is thus crucial to our understanding of how our blood performs vital systems-level functions.

Since the 1970s, mathematical models have been used to study the dynamics of RBC metabolism [42]. Other modeling formalisms, like constraint-based modeling methods [51], have been used to study mechanisms underlying cellular metabolism [8, 38, 53]. These constraint-based methods have evolved to allow for the study of system dynamics [27, 48, 9], although kinetic models are best suited to exploring temporal dynamics at short time scales [20]. From the first whole-cell kinetic model of RBC metabolism in the late 1980s [22, 23, 24, 25], researchers have examined various aspects of RBC metabolism through kinetic modeling including regulatory network structure [41], the role of 2,3-diphosphoglycerate (2,3-DPG) in binding hemoglobin [32, 34, 33], hereditary glucose-6-phosphate dehydrogenase (G6PDH) deficiency [35], and personalized pharmacodynamics [7]. There lies a gap, however, between what can be computationally modeled using kinetic approaches (limited by parameterization and network scale) and constraint-based approaches (limited by the completeness of the network structure).

Here, we address this gap with the development of a whole-cell model of the metabolism of the human Red Blood Cell and Macromolecular Mechanisms (RBC-MM). This model is built from first principles and uses kinetic and thermodynamic constraints to represent macromolecular mechanisms such as O_2_ and CO_2_ binding to hemoglobin and the detoxification of reactive oxygen species (ROS). Further, we account for the limitation to maximum reaction velocity due to finite enzyme abundance and effective turnover rate through the integration of recently published targeted proteomic data [10]. We validate our model against previous computational models and experimental data, demonstrating the ability of the RBC-MM to compute physiologically meaningful quantities and phenotypes, such as the oxygen dissociation curve (ODC). We anticipate that the computational modeling framework presented here will help usher in a new frontier in metabolic modeling that integrates non-metabolic mechanisms to better represent the functions of complex biological systems.

## RESULTS

We begin by detailing the construction and scope of the RBC-MM model. We then proceed to demonstrate its simulation capacity, providing validation of various computed phenotypes and quantities. Finally, we compute systems-level properties of the RBC network.

### Model construction

The RBC-MM model is constructed starting from the iAB-RBC-283 metabolic network reconstruction of RBC metabolism. iAB-RBC-283 represents a knowledgebase which includes 283 metabolic genes, 292 reactions, and 267 metabolites [6]. These metabolites include small molecules, cofactors, and trace minerals. For a model having *n* reactions and *m* metabolites, flux balance analysis (FBA) [39] computes the optimal metabolic state of a cell. A metabolic state is defined by a vector of reaction fluxes *v* ∈ ℝ^*n*^ (in mmol/gDW/h), by solving the linear program:

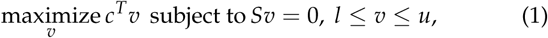

where *c* ∈ ℝ^*n*^ is the vector of objective coefficients, *S* ∈ ℝ^*m*×*n*^ is the matrix of reaction stoichiometries, and *l* ∈ ℝ^*n*^, *u* ∈ ℝ^*n*^ are the lower and upper bounds on the fluxes.

We have expanded this model to account for macromolecules present in the RBC (Figure 1). Specifically, we (*i*) account for the limitation to maximum reaction rate due to finite enzyme abundance and effective turnover rate and (*ii*) limited total protein mass in a cell. We implement these constraints in a manner similar to previous methods [3, 2]. The FBA problem associated with this expanded model is called protein-constrained FBA (P_con_FBA) [36]. In addition to fluxes, P_con_FBA also computes enzyme complex concentrations *e* ∈ ℝ^*r*^ (in mmol/gDW), and protein concentrations *p* ∈ ℝ^*s*^ (in mmol/gDW). We constrain the total protein mass (*P*) using the molecular weight of each protein (*w*). Each protein concentration is constrained by the measured concentration (Ф), while accounting for measurement error (*ϵ*). Protein concentrations are also constrained by enzyme complex stoichiometry (*a*). Given the above constraints, the P_con_FBA problem is formulated as follows:

**Fig. 1.**
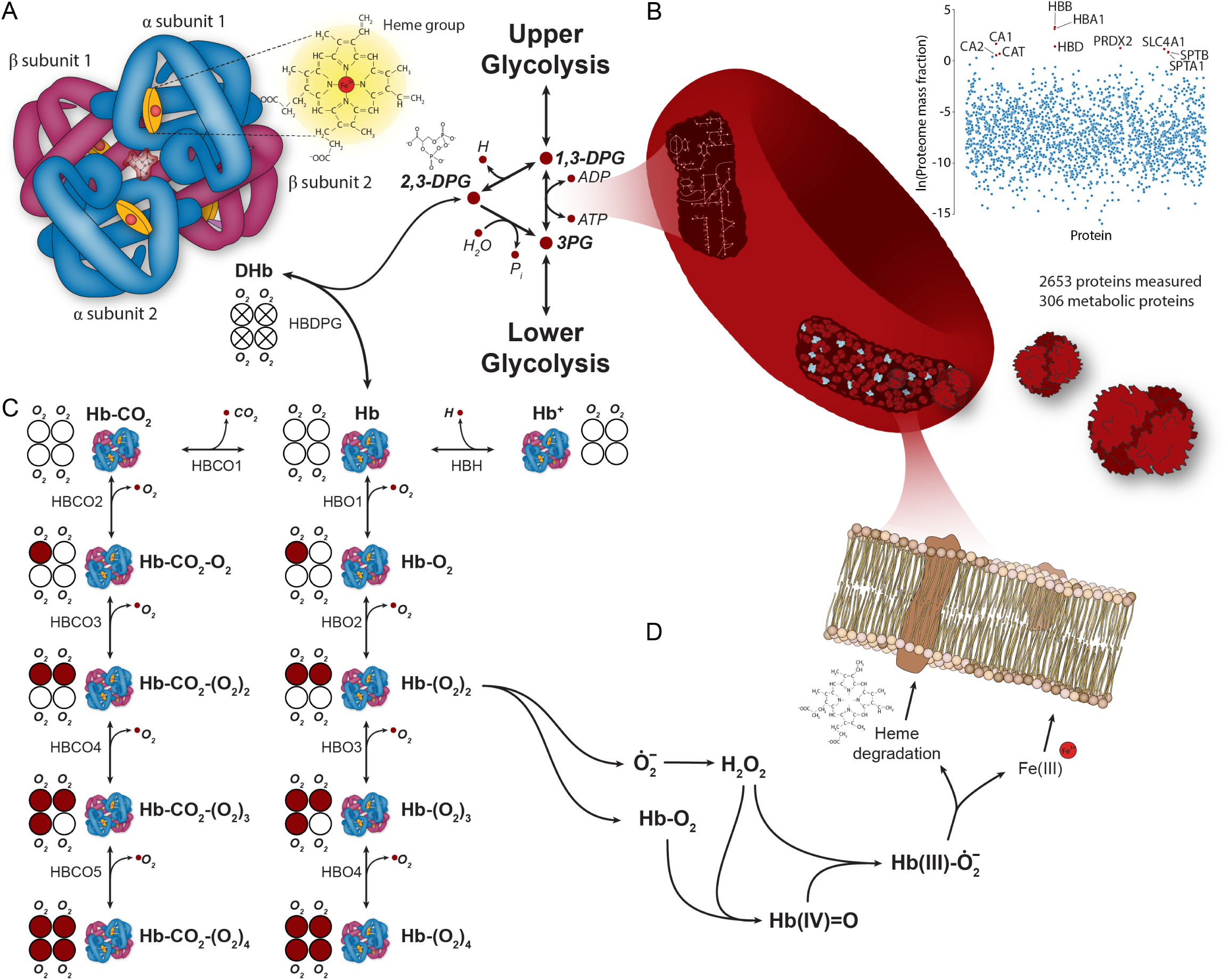
Overview of the RBC-MM model’s scope, including (**A**) integration of Hb reactions with the metabolic network, (**B**) quantitative protein constraints and crowding constraints (measured data from [10]), (**C**) Hb binding reactions, and (**D**) ROS detox mechanisms.

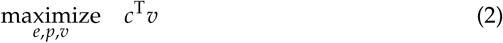

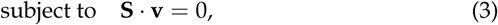

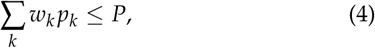

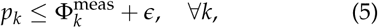

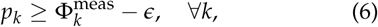

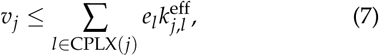

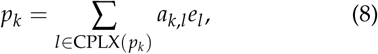

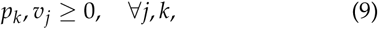

where CPLX(*j*) is the index set of reactions that are catalyzed by enzyme complex *j*, and 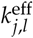 is the effective rate constant for the reaction *l* being catalyzed by enzyme *j*. The model includes quantitative constraints for 306 biochemical reactions and accounts for 2,653 total proteins using published proteomic data [10].

Next, we mathematically extend this protein-constrained model with key molecular mechanisms that describe RBC physiology.

#### Hemoglobin mechanisms

The RBC plays a vital role in the respiratory function of blood, delivering dissolved oxygen and oxygen bound to hemoglobin (oxyhemoglobin) from the lungs to the tissues [17]. In return, tissues transport CO_2_ back to the lung, where carbonic anhydrase converts biocarbonate into CO_2_ and H_2_O. The CO_2_ can be transported dissolved in the blood or can be bound to hemoglobin, depending on the concentration of O_2_ and CO_2_ in the blood [29]. The Bohr Effect [44, 4] describes how the ODC shifts based on the concentrations of CO_2_ and protons affect the affinity of hemologlobin for O_2_; the Haldane Effect describes how the ODC shifts based on the concentration of O_2_ affects the affinity of hemoglobin for CO_2_ and protons. The ODC is also dependent upon the intracellular of 2,3-diphosphoglycerate (2,3-DPG), a glycolytic intermediate and competitive inhibitor of oxygen binding to Hb [28].

We model the kinetics of hemoglobin (Hb) binding with oxygen, carbon dioxide, and 2,3-DPG. We use a cooperative mechanism (i.e., the affinity for oxygen increases with more bound oxygen) with allosteric inhibition by 2,3-DPG [40]. We model the binding of CO_2_ to Hb to account for the Bohr Effect. Here we use the notation 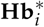 to denote a relaxed (i.e., active) form *i* of Hb and the notation **Hb**^†^ to denote a tense (i.e., inactive) form; *i* indicates the number of oxygen species bound to Hb. The full reaction schema (Figure 1C) is detailed below:

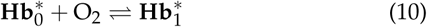

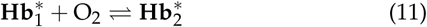

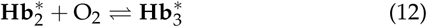

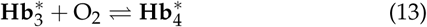

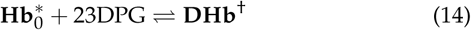

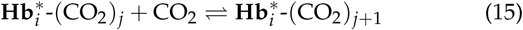

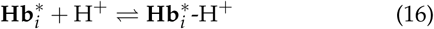

where *i* ∈ {0, 1, 2, 3, 4} represents the number of bound O_2_ molecules and *j* ∈ {0, 1, 2, 3} represents the number of bound CO_2_ molecules. To model these Hb binding kinetics, we introduce additional variables for the concentrations of Hb (all possible bound states), O_2_, CO_2_, 2,3-DPG, H_2_O, OH^−^, and H^+^.

#### Generation and detoxification of reactive oxygen species (ROS)

Hb in its alternative oxygen-bound forms is prone to auto oxidation [30], resulting in the formation of ROS. We reconstructed and included the following ROS generation reactions that involved Hb [43] (Figure 1D) in the RBC-MM model:

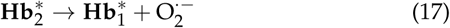

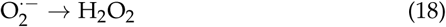

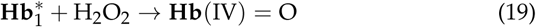

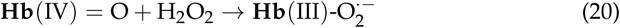

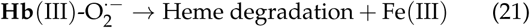

where the heme degradation product results in membrane damage, particularly observed in senescent RBCs [43]. Further, we explicitly model ROS detoxification through superoxide dismutase (Equation 22) and catalase (Equation 23):

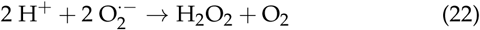

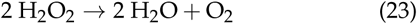

where the rate of both reactions is extremely high [1, 43].

### Model simulation

The modeling formalism described above for the RBC-MM model is built from first principles and takes a number of typically measured hematological variables as input (Table 1).

**Table 1.** Input parameters for the RBC-MM model; defaults are set according to literature values for hemoglobin concentration [12] and 2,3-DPG concentration [11].

| Input variable | Default (physiological) value | Units |
| --- | --- | --- |
| Hemoglobin concentration | 15.5 | g/dL blood |
| Hematocrit | 45 | % |
| Altitude | 0 (sea level) | m |
| Temperature | 37 (body temperature) | °C |
| Partial pressure carbon dioxide ($p\text{CO}_2$) | 40 | mmHg |
| Intracellular pH | 7.24 | - |
| 2,3-DPG concentration | 4.47 | mmol/L |

#### Computing the oxygen dissociation curve

We validated the physiological utility of the RBC-MM model by computing the oxygen dissociation curve (ODC) and comparing it to experimentally measured values. We explored three primary validations (Figure 2): (*i*) simulation of the ODC for nominal physiological values, (*ii*) qualitatively reproducing the Bohr Effect, and (*iii*) quantitatively recapitulating measured shifts due to varying concentrations of 2,3-DPG. We computed the curve using nominal baseline parameter values for the model (Table 1) according to the equation:

**Fig. 2.**
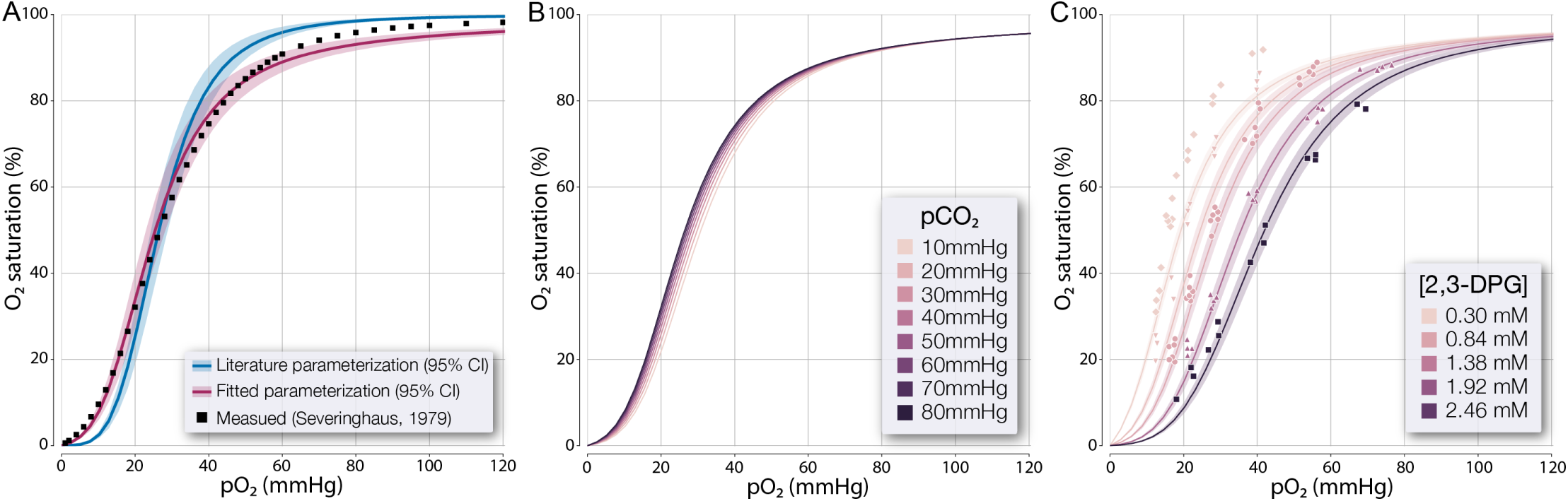
Computation of the oxygen dissociation curve (ODC). (**A**) Baseline computation for physiological values; measured data from [47]. (**B**) Computing the Bohr Effect demonstrates the expected qualitative shift of the ODC to the right with increasing pCO_2_. (**C**) Computing the effect of varying 2,3-DPG concentrations on the ODC; measured data from [16].

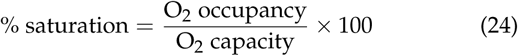

where the occupancy and capacity represent all binding states of Hb. We found that the curve is highly sensitive to the K_eq_ values used for the binding of Hb to various substrates. To examine this behavior, we used values from the literature [13, 19] and also fit the those parameters to measured data [47] (Figure 2A). It is not surprising that simulations using literature parameterization does not match measured data due to the differences between *in vitro* and *in vivo* K_eq_ values (see Supplemental Data).

Next, we tested the model’s ability to qualitatively reproduce the Bohr Effect—i.e., a shift of the ODC to the left with increasing pCO_2_ (Figure 2B). We observed that the model did indeed reproduce this behavior, matching the same qualitative trends previously observed from Hill-type models [14, 13].

We then tested the model’s ability to recapitulate the well-described dependence of the ODC on the intracellular concentration of 2,3-DPG [31, 26, 16]. We parameterized the model with measured values [16] using the reported hematocrit (40%), pH (7.195), and Hb concentration (14.75 g/dL blood); the osmotic coefficient of Hb [45] was used to convert the reported Hb concentrations. We performed a sweep across the five different concentrations of 2,3-DPG reported by Duhm [16] (Figure 2C): 0.1, 1.9, 4.4, 11.6, and 23.0 (*µ*moles/g RBC). These values were converted to units of mmol/L RBC before integration with the model. The model closely predicted the ODC for higher concentration values but underpredicted the saturation at lower concentrations. This result is likely due to parameterization of the K_eq_ values.

#### Computing the concentrations of hemoglobin bound states

In the modeling formalism described here, each Hb form is represented as a separate variable. Thus, we can compute the concentration of each bound state of Hb by solving a system of linear equations based on the K_eq_ values for each binding reaction [40]. The computed concentrations of all Hb bound states for the baseline simulation with fitted K_eq_ values (Figure 2A) are reported in Table 2. There is no experimental data with which to validate these computationally predicted values, but the values are in general agreement with estimates.

**Table 2.**
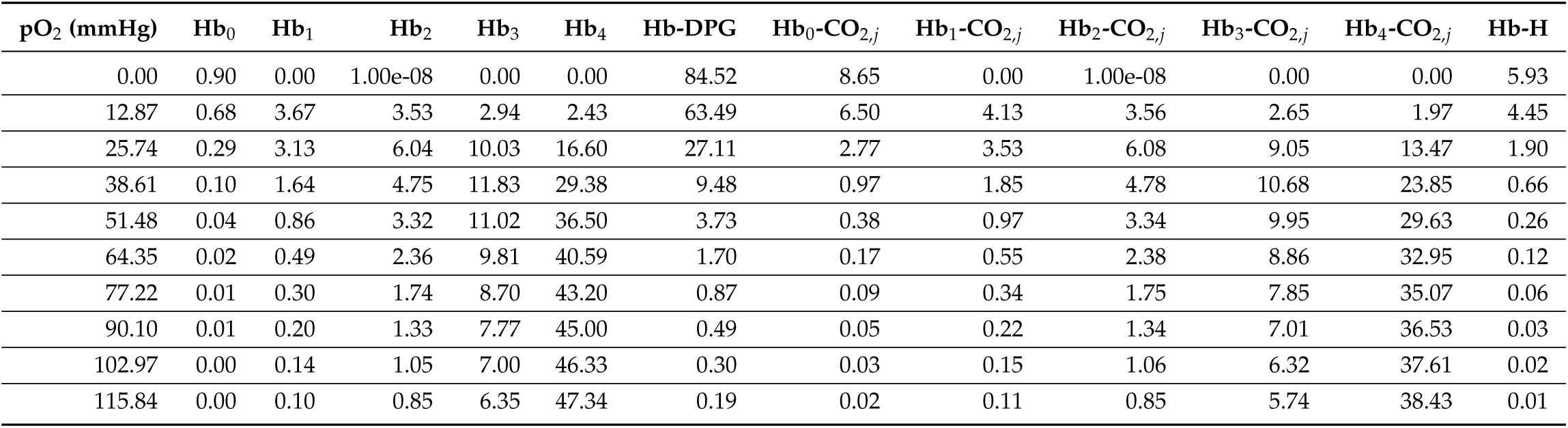
Computed fractions of hemoglobin binding states (in percentages of total hemoglobin); *j* ∈ {0, 1, 2, 3, 4} represents the number of bound CO_2_ molecules.

## DISCUSSION

The human RBC plays a crucial role in higher-level physiological function through regulating gas exchange and transport. While a variety of modeling formalisms have been used to model some of these mechanisms, the most prevalent model type to date is a Hill-type model [13], an empirical model. We have built a model that integrates these important macromolecular mechanisms with the complete known metabolic network of the RBC to compute functional states and phenotypic properties of the RBC. These results have several primary implications.

First, the modeling formalism presented here allows for the computation of physiological-scale phenotypes using mechanistic information. Such a framework will allow for the integration of whole-body deep-phenotyping data [50] to help identify causative factors important for the onset and progression of blood disorders. While constraint-based models have previously been used to explore whole-body physiology [5], these efforts were limited to studying the metabolic network. Here, we have extended the capabilities of the model to include non-metabolic macromolecular mechanisms that perform crucial physiological functions in our cell type of interest.

Second, we have computed the ODC of the human RBC from first principles using a model parameterized with standard hematological measurements. In other words, we model the binding of Hb to its various substrates using kinetic relationships instead of empirical relationships (e.g., a Hill-type model [13]). Such a modeling formalism allows for the explicit computation of the concentration of bound Hb forms. While the technology does not yet exist to experimentally measure these compounds, the ability to computationally predict them—while validating the overall phenotypic predictions—provides insight into the functional state of the blood’s gas transport and exchange system.

Third, this framework will enable the computation of personalized phenotypic responses. The use of electronic health records for personalized medical research has been increasing over the last decade [21, 49, 18], but we still need a unifying framework for the elucidation of causal biological mechanisms from such data [52]. The work presented here represents an important step in this direction, providing a computational network-based framework that integrates deep-phenotyping data typically recorded in a patient electronic health record (e.g., hematocrit, Hb concentration) to provide mechanistic predictions. By nature, these predictions offer mechanistic explanations for the observed model behavior.

Here, we have reported the most comprehensive model of the RBC to date that comprises the complete metabolic network, primary ROS detoxification, and O_2_, CO_2_, and 2,3-DPG binding to Hb. We validated this model by computing the ODC and comparing with experimentally measured values under different physiological conditions. The scope of this model could yet be expanded, namely through the addition of nitric oxide signaling, which will allow for additional physiologically relevant phenotypic predictions and insights. Taken together, this work represents a step toward marrying physiological deep-phenotyping data with a mechanistic computational framework capable of providing causal insights into the physiology of human blood.

## METHODS

To compute the ODC, we solve the optimization problem described in Equations (1-9) as we sweep across different pO_2_ values. At each pO_2_ value, we recompute the concentrations of all Hb bound states. For the baseline simulations shown in Figure 2, we selected K_eq_ values from a normal distribution using a 15% standard error 50 simulations and report the 95% confidence intervals for the 50 simulated curves, showing the mean trajectory; for the 2,3-DPG simulations, we selected K_eq_ values from a normal distribution using a 5% standard error for 50 simulations and report the 95% confidence intervals, showing the mean trajectory. The 2,3-DPG concentrations from [16] were converted into units of mmol/L RBC using the reported density of RBC as 1.110 g/mL [37].

Pseudo-first-order elementary rate constant (kPERC) values [40] are used to help sample the kinetically feasible flux solution space through setting constraints on reactions. We used literature values for “known” *k*^cat^ values and sampled remaining *k*^eff^ values from

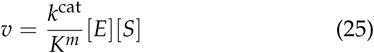

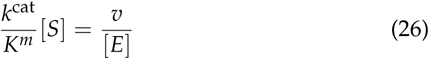

where the quantity 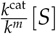 is *k*^eff^. Sampling was performed with a lower bound for the objective function, sodium-potassium transport (NaKt), set to 90% of the maximum value to estimate the physiologically relevant solution space.

The concentrations of various macromolecule bound states are modeled based on K_eq_ values where available:

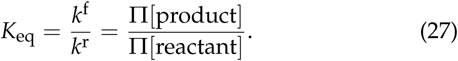

The total mass of Hb is constant,

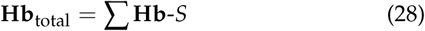

where *S* represents all bound species: O_2_, CO_2_, 2,3-DPG, and H^+^. This formulation allows us to solve the system of equations represented by Equations (27) and (28) for the concentration of each bound state as previously described [40]. See the Supplemental Methods for full details on the solving procedure and model parameterization.

## ACKNOWLEDGEMENTS

The authors would like to thank Zachary Haiman, Bin Du, Nathan Mih, Jared Broddrick, and Miguel Alcantar for valuable discussions. This study was funded by the Institute for Systems Biology’s Translational Research Fellows Program (J.T.Y.).

